# Single-chain and condensed-state behavior of hnRNPA1 from molecular simulations

**DOI:** 10.1101/2022.04.13.488036

**Authors:** D. Janka Bauer, Lukas S. Stelzl, Arash Nikoubashman

**Author notes:** Electronic mail (Arash Nikoubashman).

## Abstract

Intrinsically disordered proteins (IDPs) are essential components for the formation of membraneless organelles, which play key functional and regulatory roles within biological systems. These complex assemblies form and dissolve spontaneously over time *via* liquid-liquid phase separation of IDPs. Mutations in their amino acid sequence can alter their phase behavior, which has been linked to the emergence of severe diseases such as cancer and neurodegenerative diseases including amyotrophic lateral sclerosis. In this work, we study the conformation and phase behavior of a low-complexity domain of heterogeneous nuclear ribonucleoprotein A1 (hnRNPA1), using coarse-grained implicit solvent molecular dynamics simulations. We systematically analyze how these properties are affected by the number of aromatic residues within the examined sequences. We find a significant compaction of the chains and an increase in the critical temperature with increasing number of aromatic residues within the IDPs. Comparing single-chain and condensed state simulations, we find much more collapsed polymer conformations in the dilute systems, even at temperatures well above the estimated *θ*-temperature of the solution. These observations strongly support the hypothesis that aromatic residues play a dominant role for condensation, which is further corroborated by a detailed analysis of the intermolecular contacts, and conversely that important properties of condensates are captured in coarse-grained simulations. Interestingly, we observe density inhomogeneities within the condensates near criticality, which are driven by electrostatic interactions. Finally, we find that the relatively small fraction of hydrophobic residues in the IDPs results in interfacial tensions which are significantly lower compared to typical combinations of immiscible simple liquids.

## I. INTRODUCTION

The discovery that liquid-liquid phase separation gives rises to biomolecular condensates and membraneless organelles is revolutionizing our understanding of cell biology^1,2^ and provides new perspectives for studying self-assembly on the nanoscale.^3^ Frequently, this phase separation is driven by disordered regions of biomacromolecules or intrinsically disordered proteins (IDPs). These biomolecular condensates can be liquid and highly dynamic, but can also maturate and solidify.^2,4^ Dysregulation of liquid-liquid phase separation and formation of toxic aggregates play important roles in neurodegenerative diseases and ageing.^5–7^ In addition, the equilibrium between dilute and condensed phases may have important implications on protein function in health and disease as these two different states of matter can potentially underpin quite different biological functions.^8^ It is of particular interest to elucidate the single-chain behavior in dilute solutions and in condensates, as such knowledge could be useful for predicting, *e*.*g*., the conformation and phase behavior of IDPs based on their specific sequence. In the past few years, important progress has been made in this regard, but there are still many open questions and seemingly conflicting results:^9^ For example, high-resolution nuclear magnetic resonance experiments on dense IDP phases have revealed that local structures on the scale of individual residues can be highly similar in dilute solution and condensates,^10–12^ whereas recent electron paramagnetic resonance^13^ and single-molecule Förster resonance energy transfer^14^ experiments have observed changes in the global structures and overall extension of IDPs in dense phases compared to dilute phases.

Computer simulations are valuable tools for studying the conformation of IDPs in solution and in condensates, as they provide molecular-level insights. Further, simulations are highly suited for characterizing phase-separated biomolecular condensates, their material properties, the underlying molecular interactions and the implications of the formation of phase-separated condensates for cell biology.^7,15–20^ Atomistic simulations provide crucial information on residual structures of dilute IDPs and how chemical details modulate local and global conformations, which can be important for understanding biological function.^21–25^ However, despite encouraging progress over the past few years,^7,16,26^ such atomistically detailed simulations are still prohibitively time-consuming for studying phase separation and phase-separated condensates. Coarse-grained protein models can greatly speed up simulations, allowing for larger system sizes and longer simulation times. For instance, coarse-grained simulations at two different levels of coarse-graining highlighted contacts favouring or disfavouring the condensation of the disordered domain of TDP-43, in agreement with trends on solvation of polar and charged side chains from atomistic simulations.^7^ Further, such simplified models can help to reveal the essential elements that shape molecular systems.^17,27^ The development of coarse-grained models is a highly active field, and great strides have been made to establish transferable forcefields which can capture the sequence specific conformation and phase behavior of IDPs.^15,20,28–35^ In “sticker and spacer” models, some amino acid residues—the stickers— engage in contacts stabilizing condensates, whereas the remaining residues serve as inert spacers.^3,36–38^ These conceptually simple models are motivated by the experimental observation that favorable (noncovalent) interchain contacts are predominantly driven by only few residue types.^39^ Within this framework, the material properties of biomolecular condensates are primarily dictated by the valence and distribution of stickers, and valuable insights have been gained through it for rationalizing many important experimental trends. Given the numerous conceptual parallels between protein chains and synthetic polymers, it suggests itself to apply basic principles of (homo)polymer theory to rationalize and predict the properties of protein chains. For example, it is tempting to deduce the phase behavior of IDPs from their single-chain conformations^23,40^ or pairwise interactions at low concentrations,^23,41,42^ but it is unclear whether such relations are generally applicable to protein chains due to their heterogeneous nature.

In this work, we study to which extent concepts from homopolymer theory can be applied to protein chains, how condensation affects the conformations of disordered proteins, and whether sticker-spacer behavior emerges in simulations with a transferable coarse-grained model. The low-complexity domain of heterogeneous nuclear ribonucleoprotein A1 (hnRNPA1) (A1-LCD) is a well characterized model system which provides an ideal test ground to benchmark and validate computational approaches, and to elucidate how sequence characteristics determine dilute and dense phase behavior.^20,43^ Recently, Martin *et al*. investigated the single-chain conformation and phase behavior of A1-LCD through both experiments and simulations.^44^ They studied the wild-type as well as several variants, finding more compact polymer conformations for sequences containing more aromatic residues. Further, the critical temperature of the solutions increased significantly with increasing number of aromatic residues. These observations were rationalized *via* Monte Carlo simulations of a sticker-spacer lattice model, in which the aromatic Phenylalanine (F) and Tyrosine (Y) residues were assigned as stickers. To better understand the role of aromatic residues for the phase behavior of protein chains, we performed molecular simulations using a protein-agnostic coarse-grained IDP model,^15^ and compared the resulting phase behavior directly to experimental data. The residue-level coarsegrained model which we employed may help to better understand the context-dependent interactions of amino acids such as Arginine (R), which is known to be a sequence-specific sticker.^45^ The net positive charge of A1-LCD chains also motivated us to understand the effect of charge-charge interactions in A1-LCD condensates, which is possible in the employed coarse-grained model.^15^ Furthermore, we characterized the chain conformations in both the dilute and dense phases and identified the relevant contacts driving phase separation.

## II. MODEL AND METHODS

We model the A1-LCD protein chains using the coarsegrained hydrophobicity scale (HPS) model developed by Dignon *et al*.,^15^ where each amino acid is represented as one spherical bead of diameter *σ*_*i*_ and mass *m*_*i*_. At this level of coarse-graining, each A1-LCD chain consists of *N* = 137 particles. The solvent is modeled implicitly, and the hydrophobicity of the various residues is included in the effective monomer-monomer pair interaction

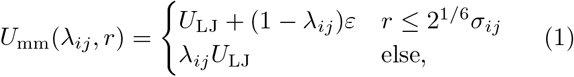

where *U*_LJ_ is the standard Lennard-Jones (LJ) potential

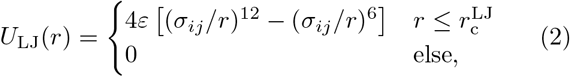

with interaction strength *ε* = 0.2 kcal/mol and cutoff radius 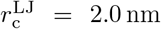^15^ The dimensionless parameter 0 ≤ λ_*ij*_ ≤ 1 controls the attraction between a pair of beads of type *i* and *j*, and thus their effective hydrophobicity. Choosing λ_*ij*_ = 0 for all pairs renders *U*_mm_ purely repulsive, resulting in good solvent conditions for the polymer. Note that bonded monomers were excluded from the pair interactions. Arithmetic averages are used as the mixing rule for the bead diameters and hydrophobicity scale, *i*.*e*., *σ*_*ij*_ = (*σ*_*i*_ + *σ*_*j*_)/2 and λ_*ij*_ = (λ_*i*_ + λ_*j*_)/2, respectively. The bead diameters *σ*_*i*_ have been set according to the van der Waals volume of the corresponding amino acids, while the λ_*i*_ are based on the hydrophobicity scale developed by Kapcha and Rossky.^46^

The employed HPS model also includes a simplified treatment of (screened) electrostatics through the pair potential

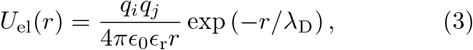

with charges *q*_*i*_ and *q*_*j*_, Debye screening length λ_D_, vacuum permittivity *ϵ*_0_, and relative dielectric constant *ϵ*_r_. To reproduce close to physiological conditions (100 mmol/L salt concentration in water), we chose λ_D_ = 1 nm and *ϵ*_r_ = 80. Further, the potential *U*_el_ has been truncated at 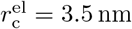 .

Bonded interactions are modeled using a harmonic potential

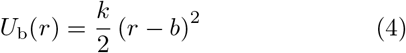

with spring constant *k* = 2000 kcal/(mol nm^2^) and equilibrium bond length *b* = 0.38 nm.

Molecular dynamics (MD) simulations were performed in the canonical ensemble, where the temperature *T* was controlled *via* a Langevin thermostat. The equations of motion were integrated using a velocity Verlet scheme with time step Δ*t* = 10 fs. All simulations were conducted on graphics processing units using the HOOMD-blue software package^47^ (v. 2.9.6) extended with azplugins (v. 0.10.2).^48^ All snapshots rendered using Visual Molecular Dynamics (version 1.9.3).^49^

## III. RESULTS

In this work, we focus on the wild-type (WT) of A1-LCD and three of its variants with fewer (Aro^−^ and Aro^−−^) and more (Aro^+^) aromatic residues (the employed amino acid sequences are identical to the ones from Ref. 44 and are included in the Supporting Information). To establish reference points, we performed simulations of freely-jointed ideal chains, achieved by disabling all non-bonded interactions and using an (effective) bond length of *b*_eff_ = 0.55 nm (see Sec. III A for details). We also simulated self-avoiding heteropolymers with the same excluded volume as the A1-LCD variants, which we realized by setting *q*_*i*_ = 0 and λ_*i*_ = 0 for all monomers *i*.

The conformation and structure formation of amphiphilic heteropolymers, such as protein chains, depend not only on the fraction of hydrophobic residues, *f*, but also on their location in the sequence.^50–52^ For example, diblock amphiphiles with *f* = 0.6 self-assemble into spherical micelles, whereas chains with uniformly distributed hydrophobic monomers merge into liquid droplets at the same *f* .^50^ To characterize the fraction and distribution of hydrophobic residues in the investigated protein chains, we calculated the sequence hydropathy decoration (SHD) parameter for each variant^51^

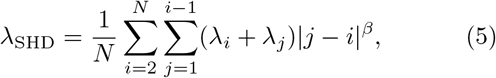

with adjustable parameter *β* = − 1.^51^ In general, λ_SHD_ increases with increasing mean hydrophobicity, ⟨ λ ⟩, and when the hydrophobic residues are spatially clustered in a sequence. Indeed, we found that both λ_SHD_ and ⟨ λ ⟩ increased monotonically with increasing number of aromatic residues (Table I). Note that the spatial distribution of charged residues is identical for all variants with a patterning parameter^21^ of 0.215, and that all protein chains have a net positive charge of 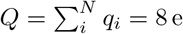 .

**Table I.** Sequence hydropathy decoration parameter λ_SHD_, mean hydrophobicity ⟨ λ ⟩, and molecular mass *M* of the four investigated variants of A1-LCD.

| Variant | $\lambda_{\text{SHD}}$ | $\langle \lambda \rangle$ | $M$ [amu] |
| --- | --- | --- | --- |
| Aro<sup>+</sup> | 5.58 | 0.62 | 13443 |
| WT | 5.48 | 0.61 | 13081 |
| Aro<sup>-</sup> | 5.33 | 0.59 | 12688 |
| Aro<sup>--</sup> | 5.18 | 0.57 | 12219 |

### A. Chain conformations

Although IDPs lack well-defined folded structures, they are characterized by a wide range of (self-similar) conformations, which strongly depend on their environment. For example, protein chains containing a large fraction of hydrophobic residues collapse into compact globules to minimize the interfacial area with the surrounding poor solvent, whereas predominantly hydrophilic chains typically display swollen coil-like conformations. In a *θ*-solvent (or at the *θ*-temperature, *T*_*θ*_, equivalently), the attraction between monomers exactly cancels the polymer-swelling due to excluded volume and charge effects, resulting in (nearly) ideal conformations.^53^ Since the intramolecular interactions depend so strongly on solvent quality, it seems likely that the interactions between different polymers, and thus their phase behavior, should follow similar trends. Indeed, for homopolymer solutions, *T*_*θ*_ becomes equal to the critical temperature *T*_c_ in the limit *N* → *∞* .^54^ Thus, it is tempting to estimate the phase boundaries of IDP solutions by determining *T*_*θ*_ from single-chain measurements.^23,40,55^ In a recent simulation study, Dignon *et al*. found a strong correlation between *T*_*θ*_ and *T*_c_ for selected IDPs,^23^ but it remains unclear how reliable such relations are for making *quantitative* predictions for the phase behavior of protein chains. For example, they found *T*_c_ ≈ 359 K and *T*_θ_ ≈ 333 K for fused in sarcoma (FUS) proteins,^23^ although one would expect *T*_*θ*_ *> T*_c_ for polymers with an upper critical solution temperature.^53,56^

To determine the conformations of the A1-LCD variants, we performed simulations in dilute as well as concentrated solutions, and computed the radius of gyration tensor for each chain

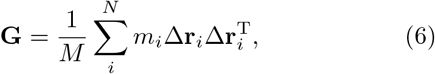

where *M* is the total mass of a protein chain, and Δ**r**_*i*_ is the position of monomer *i* relative to the polymer’s center-of-mass. The root mean square radius of gyration was taken as *R*_g_ ≡ ⟨ *G*_*xx*_ + *G*_*yy*_ + *G*_*zz*_ ⟩ ^1/2^, which is shown in Fig. 1(c) at various temperatures. At infinite dilution, *R*_g_ increased monotonically with increasing temperature, indicating an improvement of the (effective) solvent quality. Further, IDPs with more aromatic residues had a smaller *R*_g_ at any given temperature, indicating a stronger hydrophobicity of those variants, consistent with their values of λ_SHD_ and ⟨ *λ* ⟩ (Table I). This behavior is in semi-quantitative agreement with recent small-angle X-ray measurements of the investigated A1-LCD variants,^44^ which established a decrease from *R*_g_ ≈ 2.9 nm (Aro^−−^) to *R*_g_ ≈ 2.4 nm (Aro^+^) as the number of aromatic residues in the A1-LCD sequence was increased at *T* = 296 K. Note that even at the highest investigated temperature, the protein chains are much more compact than a self-avoiding heteropolymer of the same contour length, which has *R*_g_ ≈ 4.1 nm according to our simulations. Recent simulations by Tesei *et al*.^20^ and Martin *et al*.^35^ showed that the original parameterization of the HPS model^15^ caused conformations which were too collapsed for some IDP sequences, and we therefore ran additional simulations of the A1-LCD (WT) sequence at infinite dilution using their re-parameterized model [black stars in Fig. 1(c)].^20^ However, the differences between *R*_g_ from the two models are negligible for A1-LCD (WT). It is tempting to determine *T*_*θ*_ for the A1-LCD variants by comparing the *R*_g_ data from our simulations to the theoretical value of an ideal chain.^51,55,57^ For a linear (self-avoiding) homopolymer, the radius of gyration is theoretically given by^58^

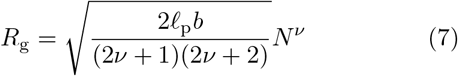

with scaling exponent *ν* = 1/3 for a collapsed globule in a poor solvent, *ν* = 1/2 for ideal-chain conformations at *θ*-conditions, and *ν ≈* 3/5 for swollen coils in a good solvent. The persistence length *𝓁*_p_ describes the local bending rigidity of the chains,^53,59,60^ with *𝓁*_p_ *≥ b/*2 even without an explicit bending potential (as is the case in the model employed here), due to local excluded volume effects. Often, *b* and *𝓁*_p_ are combined into an effective bond length 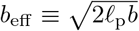, so that the radius of gyration of an ideal chain can be written as *R*_g,id_ = *b*_eff_ (*N/*6)^1/2^, with *b*_eff_ = *b* for a freely-jointed ideal chain. For self-avoiding homopolymers, *b*_eff_ and *ν* can be determined by calculating *R*_g_ for a series of different *N* and then fitting the results to Eq. (7). Alternatively, *b*_eff_ and *ν* could be determined at one fixed *N* by calculating *𝓁*_p_ from the orientational correlation of the bond vectors,^53^ and then varying *ν* in Eq. (7) until the theoretically predicted *R*_g_ matches the measurement.

**Figure 1.**
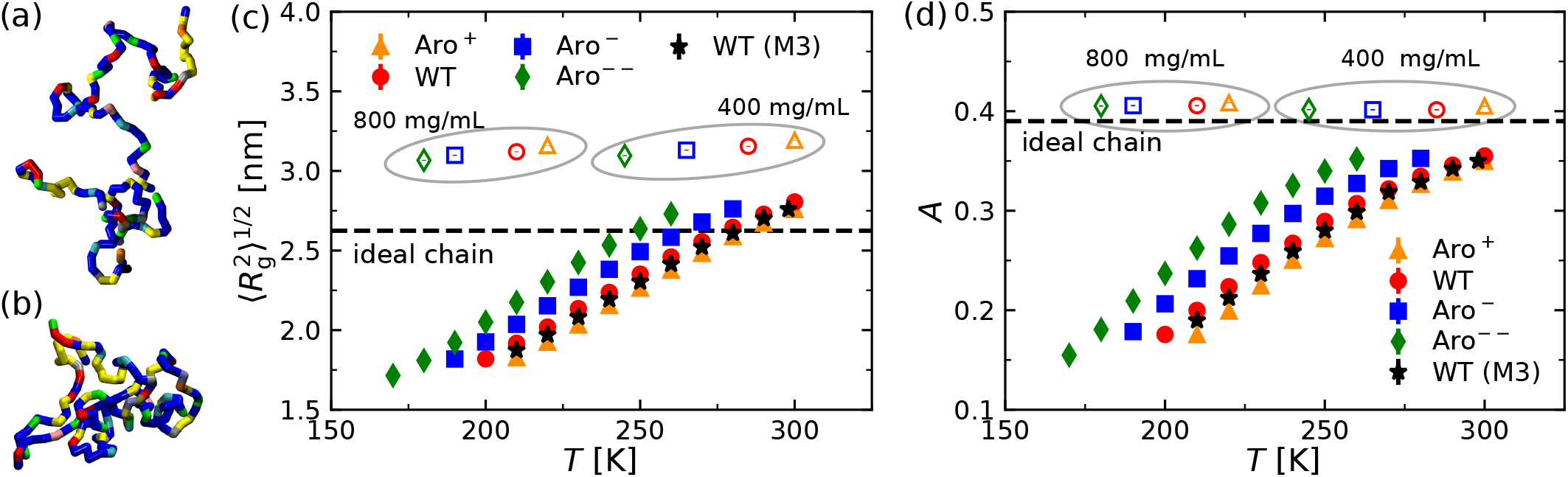
(a, b) Simulation snapshots of A1-LCD (WT) chains in dilute solution at (a) *T* = 300 K and (b) *T* = 200 K. Colors indicate the different amino acid types. (c) Root mean square radius of gyration *R*_g_ and (d) relative shape asymmetry parameter *A* of the simulated A1-LCD variants as functions of temperature *T* . In both panels, filled and open symbols show results from single chain and bulk simulations, respectively. The data indicated by black stars have been computed from simulations using model M3 from Ref. 20. The horizontal dashed lines in (c) and (d) indicate the values of *R*_g_ and *A* for an ideal chain. Error bars are smaller than the symbol size.

It is, however, not *a priori* clear whether these approaches can be generally applied to protein chains because of their inherent heterogeneity: Given the strong sequence-dependence of heteropolymer conformations,^52^ it is unclear how chains with different numbers of residues *N* but identical scaling behavior should be constructed for protein chains without repeating patterns. Moreover, using Eq. (7) assumes that the chain conformation can be described using a single, chain-averaged *b*_eff_, which seems rather unlikely given the large differences in monomer diameters and interactions. Despite these caveats, in previous works^51,55,57^ *θ*-conditions were often identified by matching the measured *R*_g_ to Eq. (7) with *ν* = 1/2 and *b*_eff_ = 0.55 ± 0.05 nm. If we follow this procedure, we find *R*_g,id_ = 2.63 nm, and thus expect *T*_*θ*_ ≈ 277 K for A1-LCD (WT). This estimate of *T*_*θ*_ is, however, unreliable as one would expect that the protein chains follow ideal chain statistics at *T* ≈ *T*_*θ*_ *independent* of polymer concentration,^53^ which is not the case here [Fig. 1(c)]. Further, we expect from Flory-Huggins theory that *T*_*θ*_ *> T*_c_ for a polymer solution with an upper critical solution temperature,^53,56^ but this condition is not fulfilled either (*T*_c_ ≈ 291 K for the wild-type, see Sec. III B).

We also characterized the chain conformations in highly concentrated solutions (*ρ*_m_ = 400 mg/mL and 800 mg/mL) at the corresponding phase coexistence temperatures (see Sec. III B). These systems contain *N*_c_ = 110 chains in a cubic simulation box with periodic boundary conditions. Figure 1(c) shows the *R*_g_ values of these systems, which are consistently *larger* than *R*_g,id_ and *R*_g_ of the corresponding protein chains at infinite dilution at the same temperature *T* . This behavior differs strongly from that of homopolymers, where one expects *R*_g_ → *R*_g,id_ with increasing polymer concentration, either by expansion of collapsed globules (poor solvents) or by the contraction of swollen coils (good solvents) due to the screening of excluded volume interactions.^53,58^ These fundamental differences ultimately stem from the fact that Flory’s ideality hypothesis^58^ cannot be applied to heteropolymers (see also the analysis of intermolecular contacts in Sec. III C).

To quantify the shape of the protein chains, we computed their relative shape asymmetry^61^

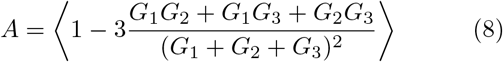

where *G*_1_, *G*_2_, and *G*_3_ are the three eigenvalues of **G**. The parameter *A* is bounded between 0 and 1, which correspond to a perfect sphere and an infinitely thin rod, respectively. The asymmetry of a three-dimensional random walk with *N* → *∞* has been determined numerically as *A* = 0.39 ± 0.004,^61^ which reflects its strong conformational fluctuations. The protein chains are much more spherical at infinite dilution compared to an ideal chain [Fig. 1(d)], likely due to the intramolecular attraction between the hydrophobic residues of the protein chains. This hypothesis is corroborated by the fact that A1-LCD variants with more aromatic residues have distinctly smaller *A* values at a given temperature *T*, and *A* approaches the value of an ideal chain with increasing *T* . In concentrated solutions, the shape asymmetry of the protein chains is *A ≈* 0.40 for all investigated systems [Fig. 1(d)], indicating that the chains have the shape asymmetry of a random walk.

To investigate the conformation of the IDPs in more detail, we computed the probability distributions *P* of the root mean square end-to-end distance, 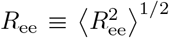, which are shown in Fig. 2 for the single chain simulations of the wild-type. The probability distributions broaden with increasing temperature, while the position of their maximum shifts to larger *R*_ee_. We have also calculated *P* (*R*_ee,id_) for a freely-jointed ideal chain with *b*_eff_ = 0.55 nm to establish a reference point (Fig. 2).^53^ The *P* (*R*_ee,id_) curve lies near the simulation results of the wild-type at *T* = 280 K, which is close to our estimate of *T*_*θ*_ based on *R*_g_. Further, the *P* (*R*_ee_) distributions of A1-LCD are much more narrow at all investigated temperatures than *P* (*R*_ee_) of a self-avoiding heteropolymer.

**Figure 2.**
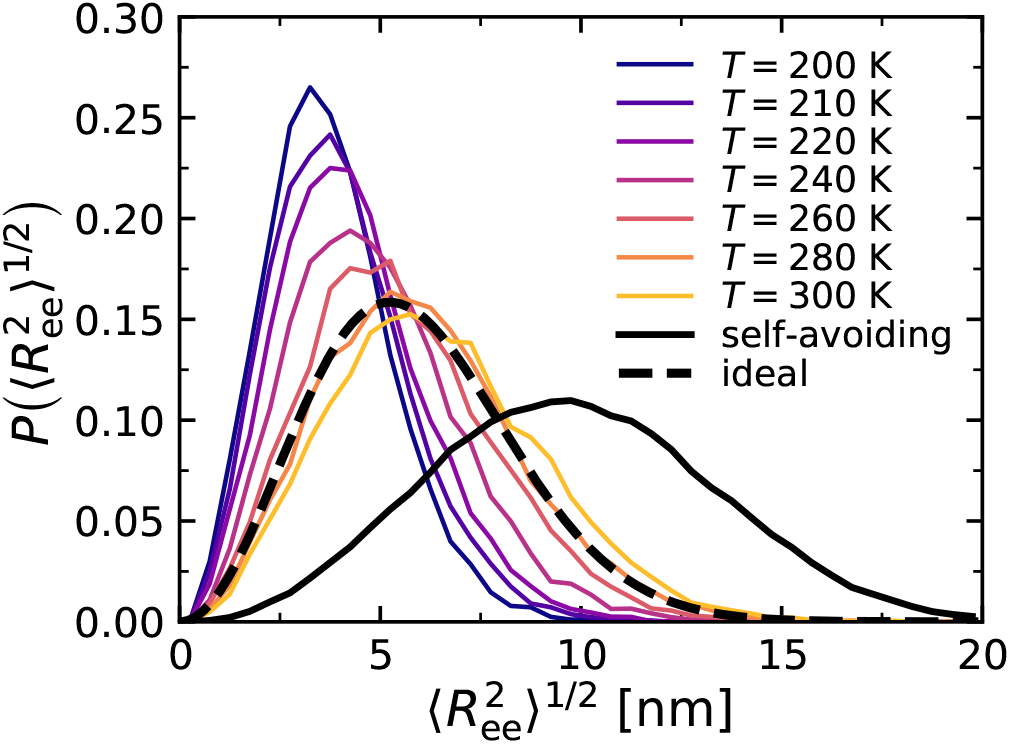
Probability distribution *P* of the end-to-end distance, *R*_ee_, from single chain simulations of A1-LCD (WT) for different temperatures (solid colored lines). Data for an ideal^53^ and a self-avoiding chain are included as dashed and solid black lines, respectively.

We have also calculated the mean interresidue distances, ⟨ (**r**_*i*_ − **r**_*j*_)^2^ ⟩^1/2^, which are shown in Fig. 3 for A1-LCD (WT) at various temperatures. These data show that the protein chains at low temperatures are locally more compact than ideal chains due to intermolecular attraction of oppositely charged and hydrophobic residues (see Sec. III C for details). At *T* = 280 K, the protein chains have similar mean interresidue distances as an ideal chain, but are much more compact than a self-avoiding heteropolymer in a good solvent.

**Figure 3.**
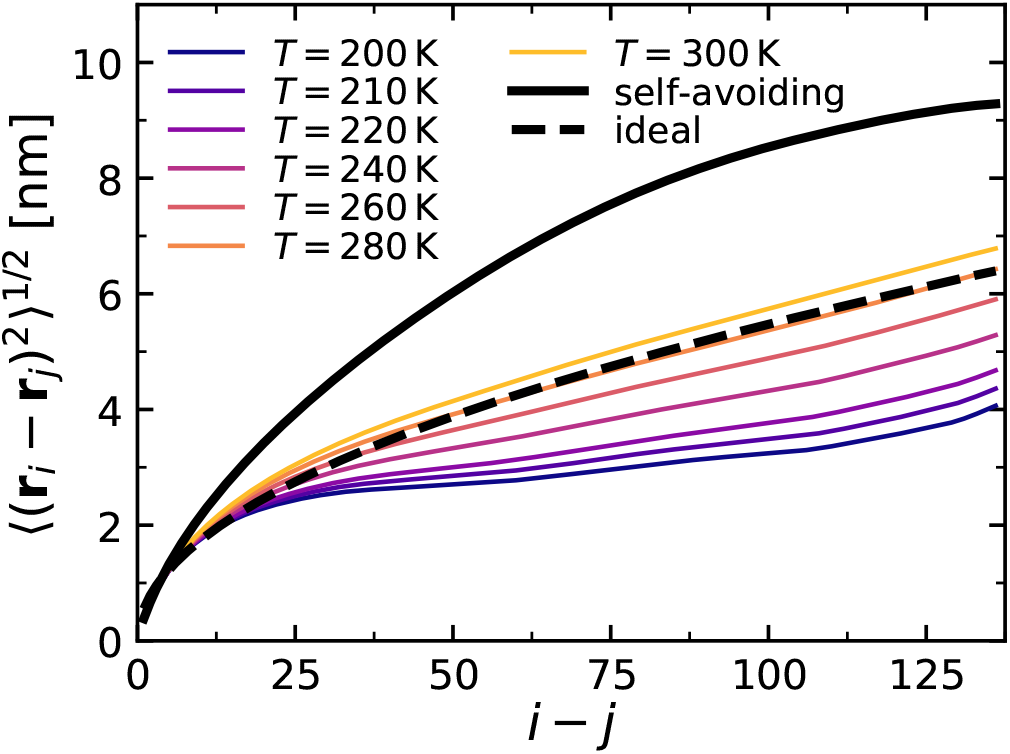
Mean interresidue distance ⟨ (**r**_*i*_ *–* **r**_*j*_)^2^⟩ ^1/2^ from single chain simulations of A1-LCD (WT) at various temperatures. Data for an ideal and a self-avoiding chain are included as dashed and solid black lines, respectively.

Although a temperature seemingly exists where certain structural properties of the A1-LCD protein chains follow ideal chain statistics, *e*.*g*., the probability distribution of the root mean square end-to-end distance, other important relations are not fulfilled at that state, *e*.*g*., the radius of gyration strongly differs in dilute and concentrated solutions. These differences are ultimately rooted in the heterogeneous nature of protein chains, which makes it impossible to describe them—even at high degree of coarse-graining—as self-similar fractal objects with chain-averaged persistence lengths. One could also argue that *T*_*θ*_ is ill-defined for heteropolymers, as the excluded volume interactions of the different residue types disappear at different *T* . Thus, we advise caution in inferring the phase behavior of protein chains from singlechain simulations, even if this strategy works reasonably well for certain cases.^23,31,41^

### B. Phase behavior

We determined the vapor-liquid phase coexistence densities of the protein chains from MD simulations containing both a low density vapor phase and a high density liquid phase in the same (elongated) simulation box.^15,62,63^ Initially, *N*_c_ = 220 chains were placed in a densely packed slab, with its two surface normals lying parallel to the *z*-axis. The dimensions of the simulation box (*L*_*x*_ = *L*_*y*_ = 15 nm and *L*_*z*_ = 150 nm) and the total number of chains were chosen such that the two vapor-liquid interfaces of the slab can be assumed as non-interacting with each other. The systems were then simulated for 10 *µ*s. Measurements were taken once the memory of the starting configuration was completely lost (typically after *t* = 1 *µ*s), which we verified by tracking *R*_g_ as well as the densities in the liquid and vapor regions. Figures 4(a-c) show selected simulation snapshots of the A1-LCD (WT) at temperatures below and above the estimated critical temperature, *T*_c_ ≈ 291 K (see further below).

**Figure 4.**
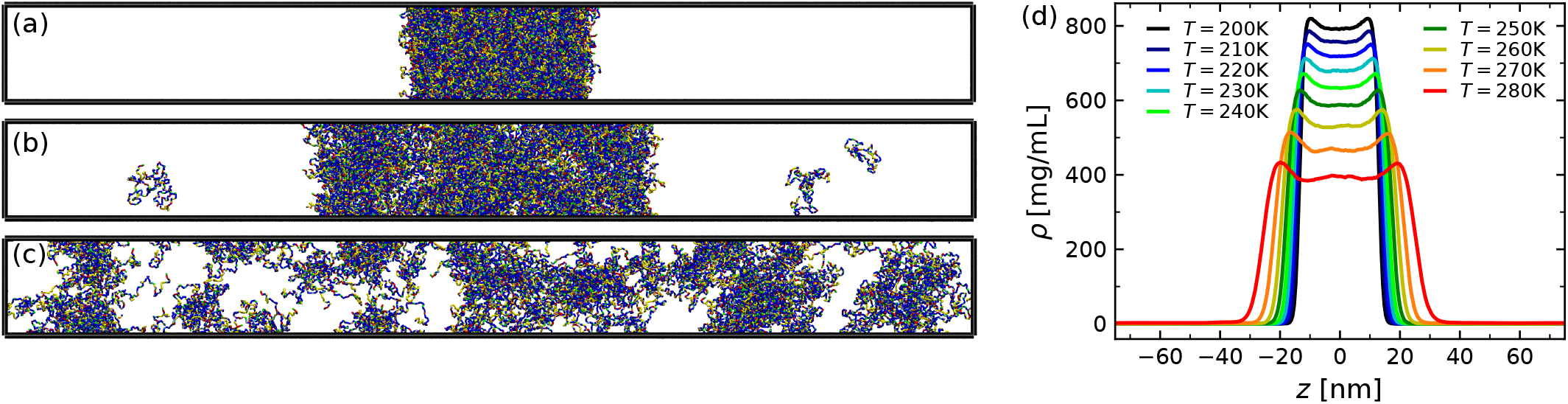
Simulation snapshots of A1-LCD (WT) at three selected temperatures (a) *T* = 200 K, (b) *T* = 280 K, and (c) *T* = 300 K. Colors indicate the different amino acid types. (d) Density profiles obtained by simulations of 220 chains of A1-LCD (WT) for various temperatures, as indicated.

Figure 4(d) shows the corresponding density profiles for various temperatures. With increasing temperature *T*, the density of the vapor phase, *ρ*_v_, increased, whereas the density of the liquid phase, *ρ*_l_, decreased, similar to the phase behavior of homopolymers in solution.^53,62,63^ Interestingly, the density profiles show microstructuring in the liquid phase, where the condensate is denser at the interface than at its center. These inhomogeneities are more pronounced at higher temperatures. The slight depletion of protein chains at the center of the condensates is also apparent in Fig. 4(b). Given the net positive charge of the A1-LCD chains (*Q* = 8 e per chain), we surmise that the intra-condensate structuring is likely due to the long-range repulsion between like-charged residues. Thus, one might expect that the positively charged residues have a tendency to face the aqueous solvent to minimize their long-range repulsion. However, we did not find such segregation, probably because the charged residues are uniformly distributed in the protein sequence. To test whether the charged residues are responsible for the microstructuring, we performed additional simulations of uncharged A1-LCD (WT) chains, and found indeed much flatter density profiles, even when the number of chains was reduced to *N*_c_ = 110 (see Fig. 5). Note also that the two-phase coexistence region extends to significantly higher temperatures when electrostatic interactions are disabled.

**Figure 5.**
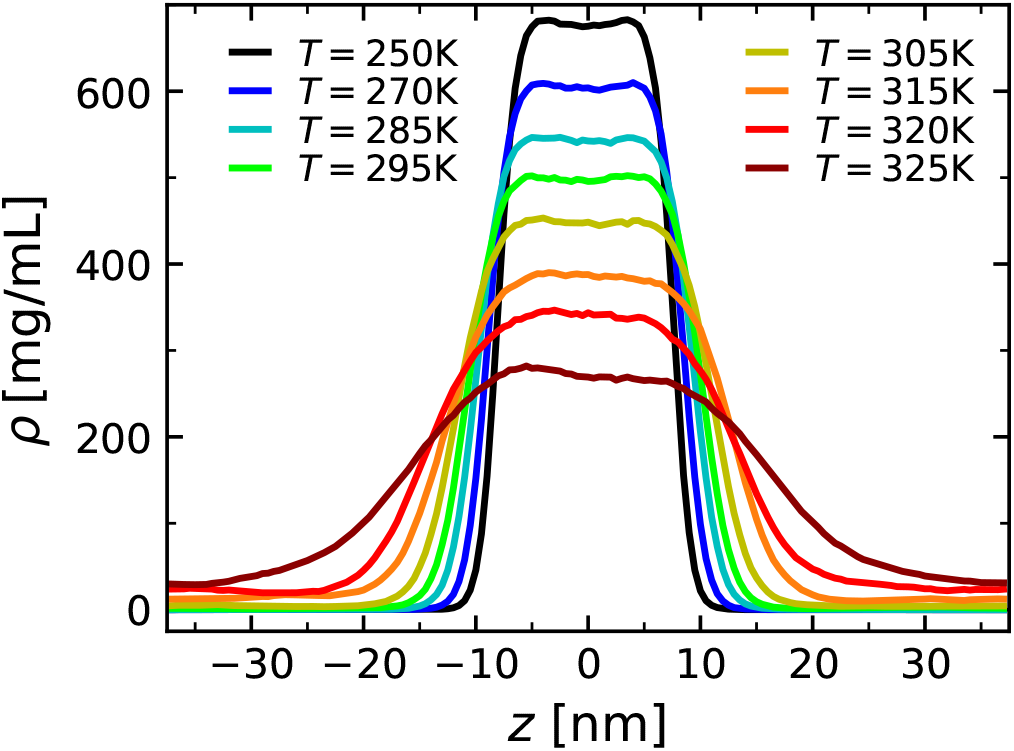
Density profiles obtained by simulations of 110 chains of A1-LCD (WT) with all charges set to zero.

We determined the critical temperature *T*_c_ by fitting the density profiles from the MD simulations to

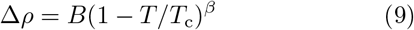

with order parameter Δ*ρ* = *ρ*_l_ *− ρ*_v_, critical amplitude *B*, and critical exponent *β* = 0.325, assuming that our systems belong to the 3*d*-Ising universality class. To estimate the critical density *ρ*_c_, we consider the rectilinear diameter *ρ*_d_

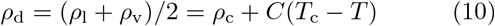

with positive (fitting) constant *C*. We also performed a finite-size scaling analysis of the critical behavior by running slab simulations with half the number of chains (*N*_c_ = 110 instead of 220), and found that the resulting systematic errors are comparable to the statistical error in determining the critical point by fitting Eq. (9).^63^

Figure 6 shows the resulting phase diagrams from simulations as well as the corresponding experimental results from Ref. 44. Based on our analysis, we find that *T*_c_ increases with increasing number of aromatic residues in the protein chains, while the critical density *ρ*_c_ remains constant within our measurement accuracy (Table II). These trends are in semi-quantitative agreement with experimental results,^44^ but *T*_c_ is about 30 − 50 K lower in the simulations compared to the experiments, while *ρ*_c_ is about 100 mg/mL larger in the simulations (see Table II). There are several potential sources for these differences, which we will briefly discuss in the following to guide future model improvements: (i) The employed model^15^ (and also its recent modification^20^) has been parameterized to match single chain conformations at infinite dilution, which does *not* guarantee correct interactions between multiple chains. Such many-body interactions could be improved through the inclusion of local-density dependent potentials.^64,65^ (ii) The IDPs are considered as fully flexible chains, and thus does not capture any secondary structures and/or local bending rigidity, which might affect the phase behavior of the protein chains.^59,60,66^ Such local chain structures could be included in coarse-grained models *via* Gō-type interactions parameterized through experimental data or atomistic simulations.^67–69^ (iii) Electrostatic interactions between charged residues are included *via* a screened Coulomb potential [Eq. (3)] with constant dielectric constant *ϵ*_r_ = 80, which is a decent approximation for dilute solutions of moderately charged protein chains in water, but less accurate for inhomogeneous dielectric media. This description of the electrostatics could be improved in the future by using, *e*.*g*., a generalized Born model.^70,71^ (iv) Both hydrophobic and electrostatic interactions are truncated at a cutoff radius, which can have a sizable impact on the phase behavior, especially near criticality.^72^ To demonstrate this effect, we ran additional simulations of A1-LCD (WT) with a smaller cutoff radius for the hydrophobic interactions (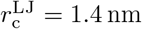 instead of 2.0 nm), finding that *T*_c_ decreased by about 60 K (see Supporting Information). Nevertheless, the agreement between simulations and experiments is rather remarkable, given the heavily coarse-grained and generic nature of the employed model.

**Figure 6.**
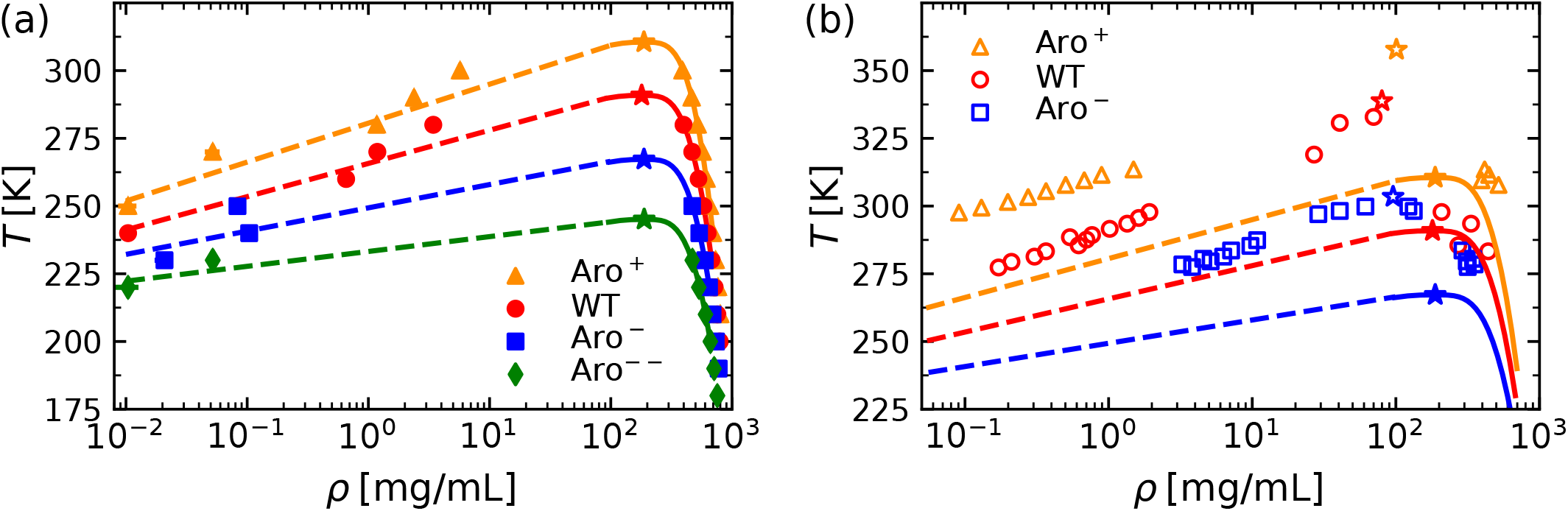
(a) Phase diagrams of the A1-LCD variants from simulations. Symbols show coexisting densities from simulations, while solid lines represent the theoretical curves, obtained by fitting the simulation data to Eqs. (9) and (10). Dashed lines are guides to the eye. (b) Comparison between simulations and experiments. Lines correspond to the fitted simulation data shown in (a), while experimental data are indicated by open symbols. Critical points are indicated by stars in both panels. Error bars are smaller than the symbol size.

**Table 2.** Critical temperatures and densities from our MD simulations (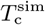 and 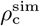) and from experiments^44^ (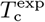 and 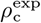) for investigated variants of A1-LCD.

| Variant | $T_c^{\text{sim}}$ [K] | $\rho_c^{\text{sim}}$ [mg/mL] | $T_c^{\text{exp}}$ [K] | $\rho_c^{\text{exp}}$ [mg/mL] |
| --- | --- | --- | --- | --- |
| Aro<sup>+</sup> | 311 ± 1 | 188 ± 6 | 357.75 | 100.9 |
| WT | 291 ± 1 | 180 ± 5 | 338.75 | 79.9 |
| Aro<sup>-</sup> | 267 ± 1 | 188 ± 6 | 303.45 | 95.6 |
| Aro<sup>--</sup> | 245 ± 2 | 189 ± 4 | - | - |

We also determined the surface tension of the protein chains, *γ*, from the slab simulations using the Kirkwood-Buff relation

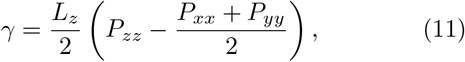

where the factor 1/2 accounts for the two interfaces present in the system. The components of the pressure tensor, **P**, are calculated *via* the standard Clausius virial equation. Note that Eq. (11) holds only for homogeneous liquid phases, and therefore we used it to compute *γ* only in the temperature regime, where the condensed phase did not show any pronounced density fluctuations. Figure 7 shows the temperature-dependence of *γ*, which goes to zero as *T* → *T*_c_, as expected. At fixed *T*, protein chains containing more aromatic residues have larger *γ* [Fig. 7(a)], which is consistent with their overall higher hydrophobicity (Table I). For biologically relevant protein concentrations in the range of *ρ* ≈ 400 − 800 mg/mL (simulated at the corresponding phase coexistence temperatures far from criticality, *cf*. Fig. 6), the computed surface tensions lied between 5 *µ*N/m and 800 *µ*N/m, which is comparable to recent experimental measurements of phase separated protein droplets.^73–75^ A similar range of *γ* ≈ 15 − 380 *µ*N/m was also measured for FUS proteins in recent MD simulations, using a very different coarse-grained model with an explicit solvent.^18^ Interestingly, these values are several orders of magnitude smaller than typical interfacial tensions of simple liquid combinations, *e*.*g*., alkane droplets in water which have *γ* ≈ 50 mN/m at room temperature.^76,77^ This large difference in *γ* persists even if one shifts the temperature in Fig. 7 to account for the mismatch of *T*_c_ between simulations and experiments (see Table II).

**Figure 7.**
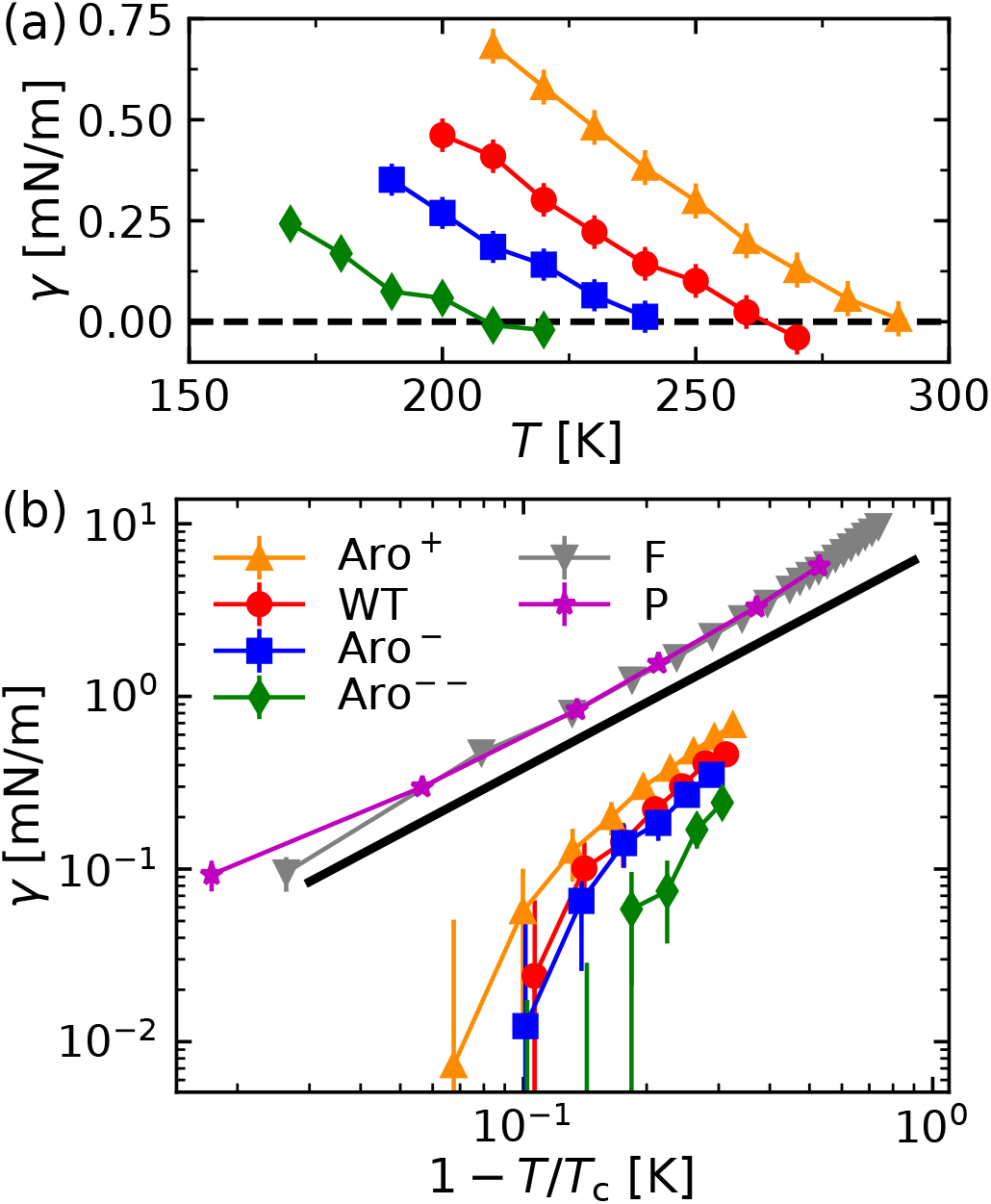
Surface tension of the protein chains as functions of (a) *T* and (b) 1 *− T/T*_c_. In panel (b), the black line indicates the theoretically expected scaling for the 3*d*-Ising universality class, *γ* ∝ (1 − *T/T*_c_) ^1.26^ .

It is known that coarse-grained models often exhibit lower surface tensions compared to experiments and atomistic models,^65,78,79^ and therefore we have performed additional simulations of highly hydrophobic Phenylalanine (F) and Proline (P) homopolymer chains (*N*_c_ = 110 in those cases) with the same degree of polymerization as A1-LCD to establish a baseline for our model. For example, we find *γ ≈* 7 mN/m for poly-F at room temperature [see Fig. 7(b) and Supporting Information for more values], which is about ten times lower than typical experimental values of alkanes in water,^76,77^ but still at least one order of magnitude larger than *γ* of the simulated A1-LCD variants in water. One might speculate that the comparatively lower surface tensions found in simulations and experiments of condensed IDPs, such as hnRNPA1 and the related protein FUS, stem from their large fraction of hydrophilic and only weakly hydrophobic residues. For instance, in the sticker and spacer representation of A1-LCD (WT) employed by Martin *et al*.,^44^ only about 15 % of residues were selected as hydrophobic stickers, which is in qualitative agreement with our contact map analysis (see Sec. III C below).

Finally, we can use the surface tension data from our simulations to verify the postulated compatibility with the 3*d*-Ising universality class, for which one expects *γ* = *γ*_0_(1 *− T/T*_c_)^*µ*^ near the critical point, with amplitude *γ*_0_ and critical exponent *µ* ≈ 1.26.^80^ We found *µ* = 1.18 ± 0.32 when fitting our simulation data in the range 1 *− T/T*_c_ ≤ 0.20, which is in decent agreement with the theoretically expected value [Fif. 7(b)]. Further, *γ*_0_ increased with increasing number of aromatic residues in the protein chains.

### C. Contact Maps

To determine the inter-chain contacts of the IDPs in the concentrated regime, we performed additional bulk simulations at *ρ* = 400 mg/mL and 800 mg/mL for all four variants (WT, Aro^−^, Aro^−−^, and Aro^+^). The systems contained *N*_c_ = 110 chains in a cubic simulation box with periodic boundary conditions, and the temperature of each simulation was set to the calculated coexistence temperature (Fig. 6). Intermolecular contacts can be identified through, *e*.*g*., distance-^15^ or energy-based^20^ metrics. Appropriate criteria can be easily defined in lattice-based simulations,^44^ due to their inherently discretized length-and energy-scales. In off-lattice simulations (and also experimental systems) there is, however, some ambiguity in defining intermolecular contacts. Typically, distance-based metrics count the number of pairs within a specified cutoff radius 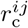 while energy-based criteria consider pairwise interaction energies.^20^ We studied the intermolecular contacts using both methods. To establish a baseline reference, we performed additional simulations of self-avoiding heteropolymers at both concentrations.

In accordance with the trends from simulations of single chains (see Sec. III A) and the phase behavior of the A1-LCD variants (see Sec. III B), we find that aromatic residues engage in many important interactions. Figure 8(a) shows the intermolecular contact map for all residues of A1-LCD (WT) at *ρ* = 400 mg/mL, using a distance-based metric where two residues are in contact when their distance is below 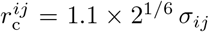 . Interchain contacts are distributed uniformly along the protein sequence, where residues that form many contacts, whether charged or aromatic, are typically followed by residues that do not engage in many contacts. To better see which residue types predominantly form contacts, we reduce the full contact map by summing over all entries with the same type of pairs, and then divide the sum by the number of possible interaction partners for each pair of types, *N*_pair_. The resulting data is shown in Fig. 8(b), highlighting that contacts between hydrophobic residues K, F, M, and Y are among the most prominent intermolecular contacts formed in the simulations of A1-LCD condensates. In contrast, A, G, and P residues have much fewer contacts although they have similar hydrophobicity parameters λ_*i*_ (see Supporting Information). Charged residues R, K, and D also form distinct contacts with their oppositely charged partners. Simulation of a purely repulsive reference system of uncharged A1-LCD (WT) chains shows that aromatic residues F and Y still form many contacts [Fig. 8(d)], which is likely due to the larger sizes of F and Y residues; this may explain why these residue types are typically more important stickers than smaller hydrophobic residues, such as A, G, and P. The comparison with the self-avoiding chain reference simulation also highlights the contacts between positively and negatively charged R-D and K-D pairs.

**Figure 8.**
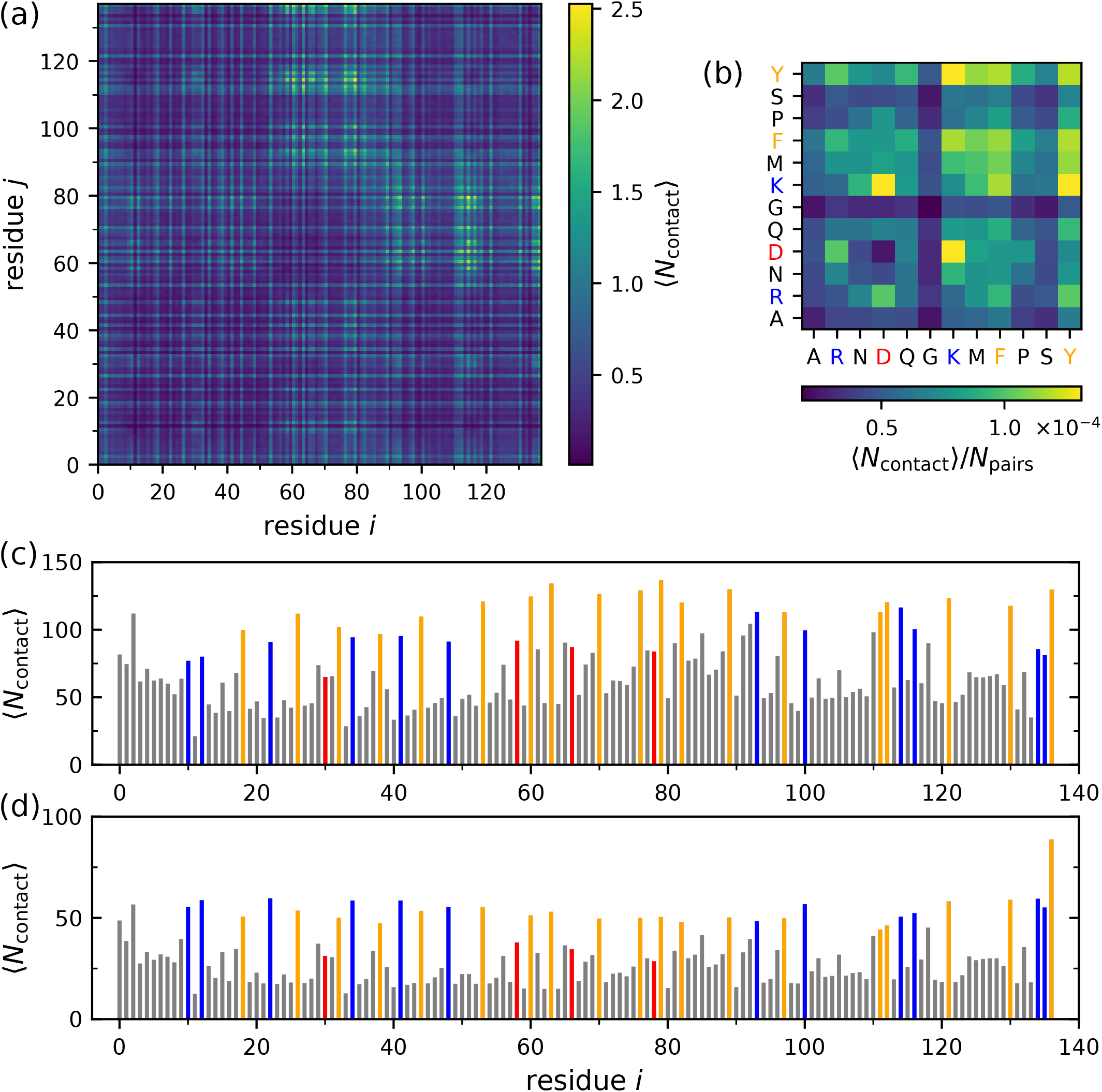
Distance-based contact maps for A1-LCD (WT) at *ρ* = 400 mg/mL and *T* = 285 K. (a) Average number of intermolecular contacts for a pair of residues *i* and *j*. (b) Average number of intermolecular contacts based on type, normalized by the corresponding number of possible interaction pairs, *N*_pairs_. (c) 1D-projection of residual contact map. (d) Same as (c) but for a heteropolymer in a good solvent. (c,d) Aromatic residues (F and Y) are marked in orange, negatively charged residues (D) are marked in red, while positively charged ones (R and K) are shown in blue.

Computing the intermolecular interaction energies provides an alternative perspective on the contact pair formation. Figures 9(a,b) show the contributions from van der Waals and electrostatic interactions, demonstrating that both aromatic and charged residues engage in stabilizing interactions, and can thus be considered as stickers. In particular, M, F, P, and Y residues are strongly attracted to each other, which reflects their high hydrophobicity.^15^ According to Fig. 9(b), R-Y interactions are favorable but less favorable than Y-Y contacts, which may require modifications of the parameter set to capture experimentally observed R-Y attraction.^32^ Considering electrostatic interactions, the negatively charged D residues are highly attracted to the positively charged R and K residues, while interactions between like-charged residues are unfavorable, as expected. Overall, electrostatic interactions are about one order of magnitude stronger than van der Waals interactions, so that two like-charged residues still repel each other even if they are highly hydrophobic. As the IDPs feature an excess of positively charged residues, it is not surprising that the negatively charged D residues engage in favorable charge-charge interactions. Recent work has highlighted that charged residues can also be important “stickers” in agreement with our results.^45^ Thus, our simulations reveal that several other residues could be regarded as stickers when constructing a sticker-spacer model of this protein chain, in addition to the aromatic F and Y residues that have been posited in recent joint experimental and simulation work.^44^ Conversely, our study in addition to other recent investigations^7,32,43^ shows that “sticker-spacer” behavior emerges naturally in coarsegrained simulations with both residue-level and multiatom resolutions.

**Figure 9.**
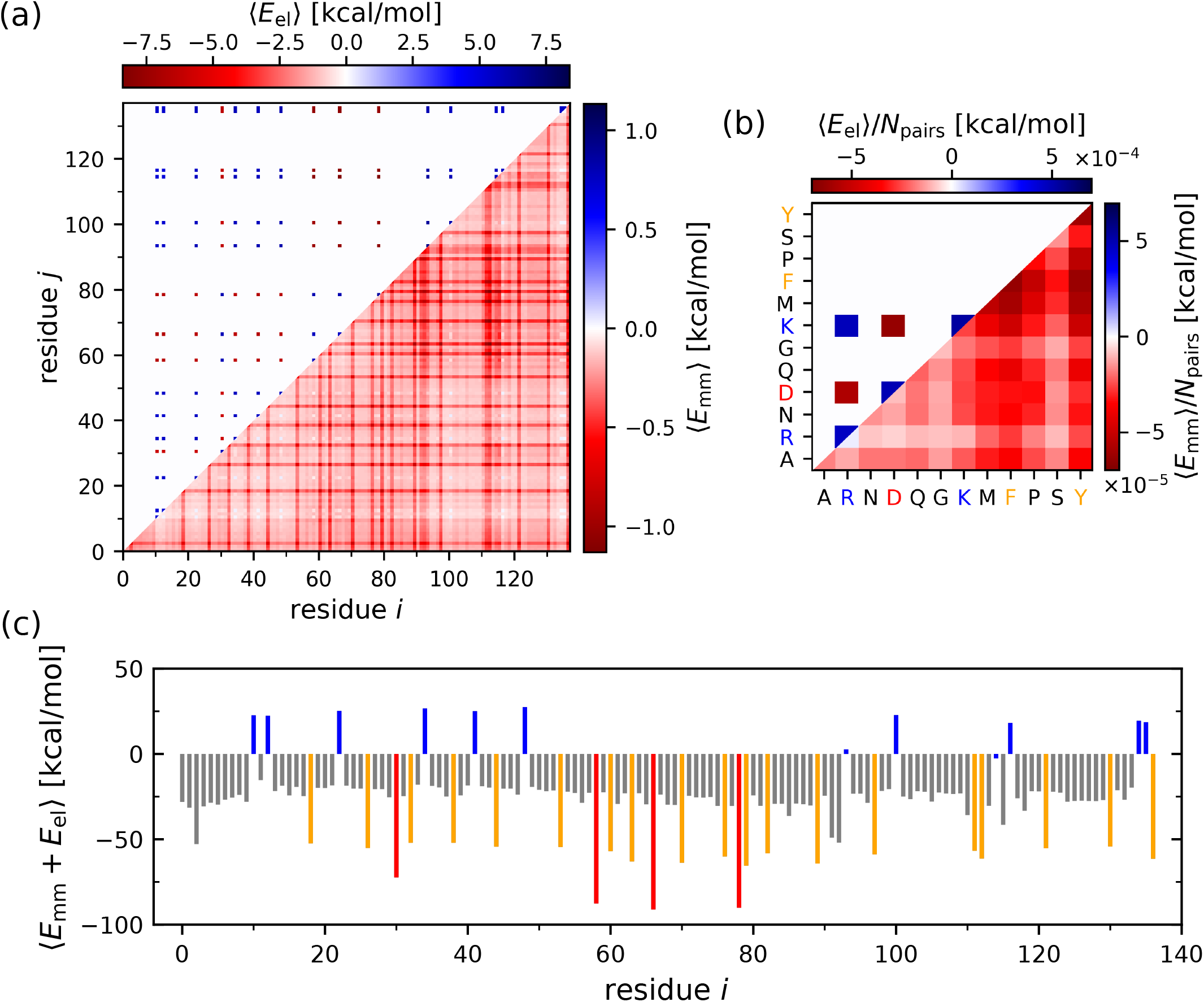
Energy-based contact maps for A1-LCD (WT) at *ρ* = 400 mg/mL and *T* = 285 K. (a) Intermolecular monomermonomer interaction energies, and (b) normalized interaction energies per type, split into electrostatic (top left half) and van der Waals interactions (bottom right half). (c) 1D-projection of interaction energies. Aromatic residues are marked in orange, charged residues are marked in red (negative) and blue (positive).

## IV. CONCLUSIONS

We have investigated the conformation and phase behavior of A1-LCD protein chains in the dilute and condensed state through molecular simulations. We investigated the wild-type as well as three mutated variants with fewer or more aromatic residues, and compared our simulations with recent experiments. Even at a relatively coarse-grained representation, with one bead per amino acid and an implicit solvent, we observed rich behavior of A1-LCD proteins and identified important trends: In dilute solutions, chains with more aromatic residues in their sequence were more compact and more spherical. In condensates, the chains were much more extended, with similar chain conformations for the different variants. These findings are particularly interesting in light of recent experiments that have also observed differences in protein chain conformations between dilute and dense phases: For example, recent single-molecule Förster resonance energy transfer experiments revealed that the intrinsically disordered protein tau, which is implicated in neurodegenerative diseases such as Alzheimer’s disease, expands in condensates.^14^ In contrast, magnetic resonance experiments found a compaction of the fused in sarcoma protein in condensates.^13^

Further, we observed that the critical temperature of the phase-separated systems increased with increasing number of aromatic residues in the sequence. To elucidate this behavior, we carried out a distance- and energy-based contact analysis, revealing that charged and aromatic residues form the majority of intermolecular contacts. We also found distinct micro-structuring in the liquid phase near criticality, driven by electrostatics, which may also occur in other condensates of charged proteins.^45^ Our simulations also revealed that the phase-separated condensates have distinctly smaller surface tensions than typical combinations of immiscible simple liquids, such as alkane droplets in water, which we attribute to the large fraction of hydrophilic residues in the protein sequences. In general, we found that the heterogeneous nature of the protein sequence, with different hydrophobic, polar, and charged amino acids, makes it difficult to apply standard homopolymer models even at the employed level of coarse-graining.

## Supporting information

Additional simulation details and results

## SUPPORTING INFORMATION

Protein sequences and additional simulation details; additional contact maps, phase diagrams, and surface tension data

## DATA AVAILABILITY

The data that support the findings of this study are available from the authors upon reasonable request.

## ACKNOWLEDGMENTS

We thank Prof. Alex S. Holehouse for fruitful discussions. This work was supported by the Deutsche Forschungsgemeinschaft (DFG, German Research Foundation) through Project Nos. 233630050, 274340645, 405552959, and 470113688. D.J.B. acknowledges support through the Max Planck Graduate Center with the Johannes Gutenberg University Mainz (MPGC). L.S.S. acknowledges support by ReALity (Resilience, Adaptation and Longevity), M^3^ODEL and Forschungsinitiative des Landes Rheinland-Pfalz. The authors gratefully acknowledge the computing time granted on the supercomputer Mogon (hpc.uni-mainz.de).

