## Additional simulation details and results for "Single-chain and condensed-state behavior of hnRNPA1 from molecular simulations"

### I. SIMULATION DETAILS

#### Studied Sequences

- A1-LCD (WT):

  GSMASASSSQ RGRSGSGNFG GGRGGGFGGN DNFGRGGNFS GRGGFGGSRG

  GGGYGGSGDG YNGFGNDGSN FGGGGSYNDF GNYNNQSSNF GPMKGGNFGG

  RSSGGSGGGG QYFAKPRNQG GYGGSSSSSS YGSGRRF

- A1-LCD (Aro$^{+}$):

  GSMAFASSFQ RGRYGSGNFG GGRGGGFGGN DNFGRGGNFS GRGGFGGSRG

  GGGYGGSGDG YNGFGNDGSN FGGGGSYNDF GNYNNQSSNF GPMKGGNFGG

  RSSGGSYGGG QYFAKPRNQG GYGGSSFSSS YGSGRRF

- A1-LCD (Aro$^{-}$):

  GSMASASSSQ RGRSGSGNSG GGRGGGFGGN DNFGRGGNSS GRGGFGGSRG

  GGGYGGSGDG YNGFGNDGSN SGGGGSSNDF GNYNNQSSNF GPMKGGNFGG

  RSSGGSGGGG QYSAKPRNQG GYGGSSSSSS SGSGRRF

- A1-LCD (Aro$^{--}$):

  GSMASASSSQ RGRSGSGNSG GGRGGGFGGN DNSGRGGNSS GRGGFGGSRG

  GGGSGGSGDG YNGSGNDGSN SGGGGSSNDF GNSNNQSSNS GPMKGGNFGG

  RSSGGSGGGG QYSAKPRNQG GSGGSSSSSS SGSGRRS

| Residue | Abrv. | Mass $m$ [amu] | Charge $q$ [e] | Diameter $\sigma$ [Å] | Hydrophobicity $\lambda$ |
|---|---|---|---|---|---|
| Alanine | A | 71.08 | 0 | 5.04 | 0.729730 |
| Arginine | R | 156.20 | 1 | 6.56 | 0.0 |
| Asparagine | N | 114.10 | 0 | 5.68 | 0.432432 |
| Aspartic acid | D | 115.10 | -1 | 5.58 | 0.378378 |
| Cysteine | C | 103.10 | 0 | 5.48 | 0.594595 |
| Glutamine | Q | 128.10 | 0 | 6.02 | 0.513514 |
| Glutamic acid | E | 129.10 | -1 | 5.92 | 0.459459 |
| Glycine | G | 57.05 | 0 | 4.50 | 0.648649 |
| Histidine | H | 137.10 | 0.5 | 6.08 | 0.513514 |
| Isoleucine | I | 113.20 | 0 | 6.18 | 0.972973 |
| Leucine | L | 113.20 | 0 | 6.18 | 0.972973 |
| Lysine | K | 128.20 | 1 | 6.36 | 0.513514 |
| Methionine | M | 131.20 | 0 | 6.18 | 0.837838 |
| Phenylalanine | F | 147.20 | 0 | 6.36 | 1.0 |
| Proline | P | 97.12 | 0 | 5.56 | 1.0 |
| Serine | S | 87.08 | 0 | 5.18 | 0.594595 |
| Threonine | T | 101.10 | 0 | 5.62 | 0.675676 |
| Tryptophan | W | 186.20 | 0 | 6.78 | 0.945946 |
| Tyrosine | Y | 163.20 | 0 | 6.46 | 0.864865 |
| Valine | V | 99.07 | 0 | 5.86 | 0.891892 |

Table S1. The 20 naturally occurring amino acids with their properties used in the HPS model.

| | $\rho = 400\,\mathrm{mg/mL}$ | | $\rho = 800\,\mathrm{mg/mL}$ | |
|---|---|---|---|---|
| Variant | Temperature $T$ [K] | Box length $L$ [nm] | Temperature $T$ [K] | Box length $L$ [nm] |
| $\mathrm{Aro}^{+}$ | 300 | 8.240 | 220 | 6.539 |
| WT | 285 | 8.163 | 210 | 6.480 |
| $\mathrm{Aro}^{-}$ | 265 | 8.082 | 190 | 6.417 |
| $\mathrm{Aro}^{--}$ | 245 | 7.083 | 180 | 6.336 |

Table S2. Temperature $T$ and box length $L = L_x = L_y = L_z$ of the bulk simulations (110 chains) for the different variants of A1-LCD.

#### II. ADDITIONAL RESULTS

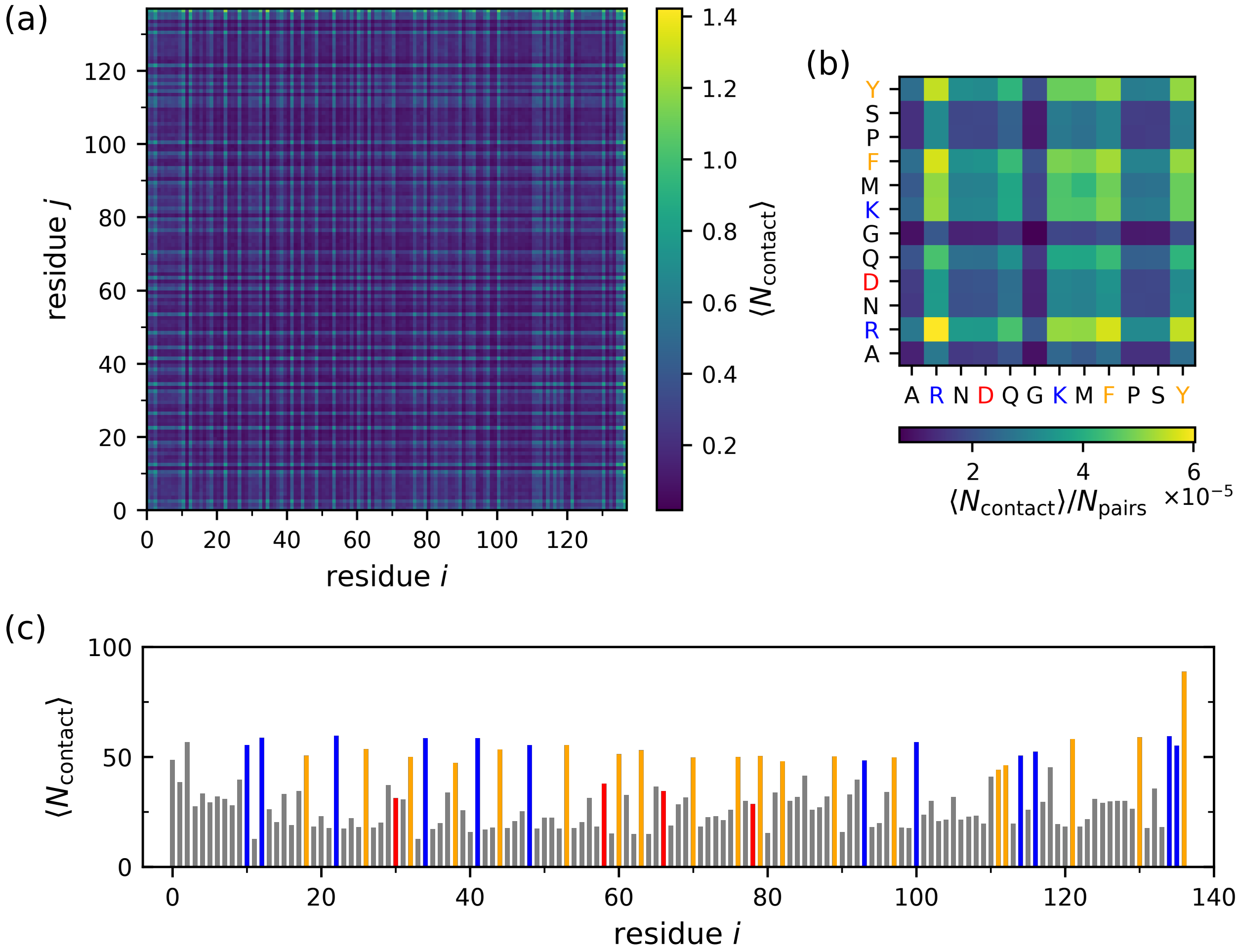


Figure S1. Distance-based contact maps for a heteropolymer in a good solvent with bead sizes according to the sequence of A1-LCD (WT). Simulations conducted at $\rho = 400\,\mathrm{mg/mL}$ and $T = 285\,\mathrm{K}$. (a) Average number of intermolecular contacts for a pair of residues $i$ and $j$. (b) Average number of intermolecular contacts based on type, normalized by the corresponding number of possible interaction pairs, $N_{\mathrm{pairs}}$. (c) 1D-projection of residual contact map. Aromatic residues (F and Y) are marked in orange, negatively charged residues (D) are marked in red, while positively charged ones (R and K) are shown in blue.

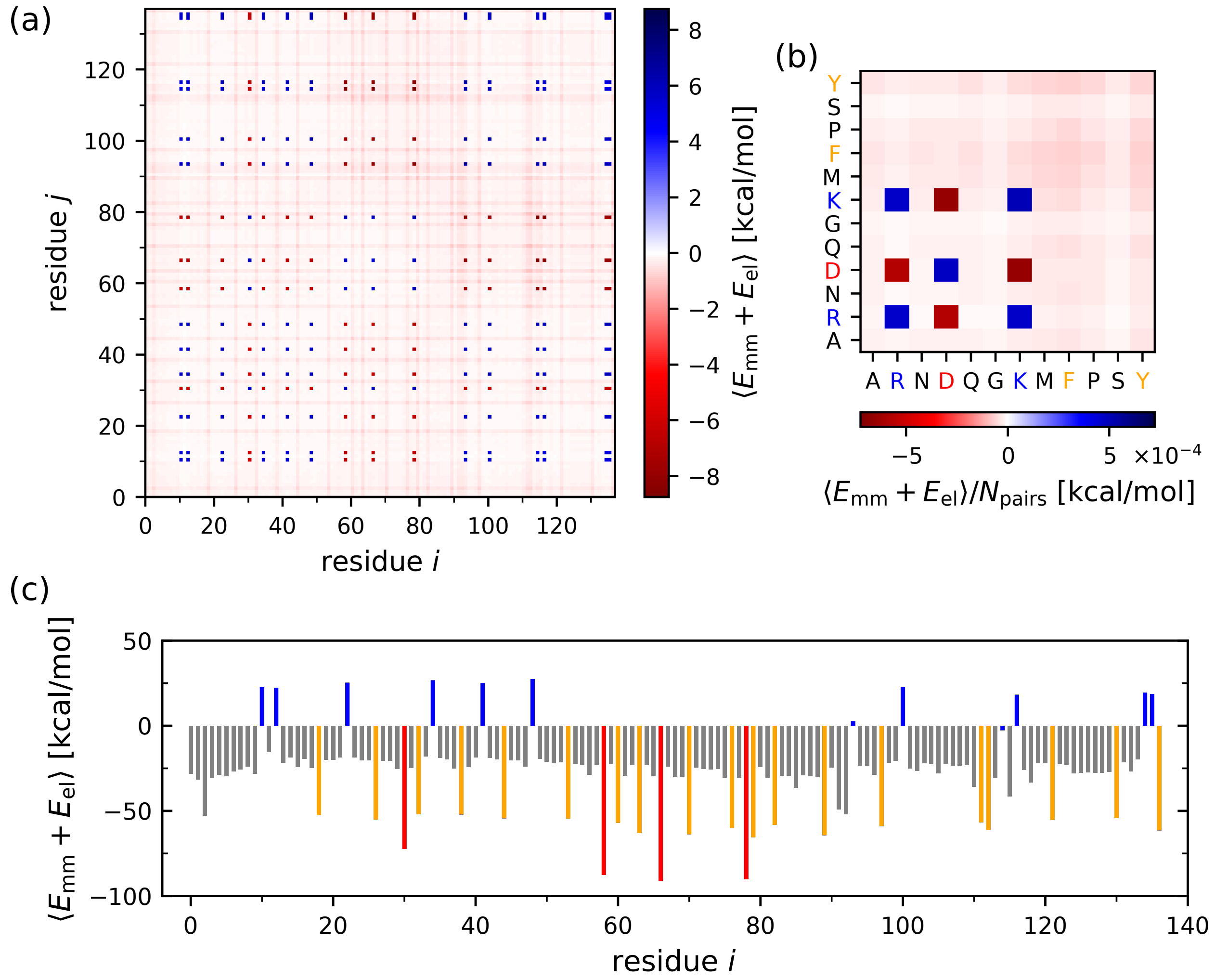


Figure S2. Energy-based contact maps for A1-LCD (WT) at $\rho = 400\,\mathrm{mg/mL}$ and $T = 285\,\mathrm{K}$. (a) Intermolecular monomer-monomer interaction energies, and (b) normalized interaction energies per type. (c) 1D-projection of interaction energies. Aromatic residues are marked in orange, charged residues are marked in red (negative) and blue (positive).

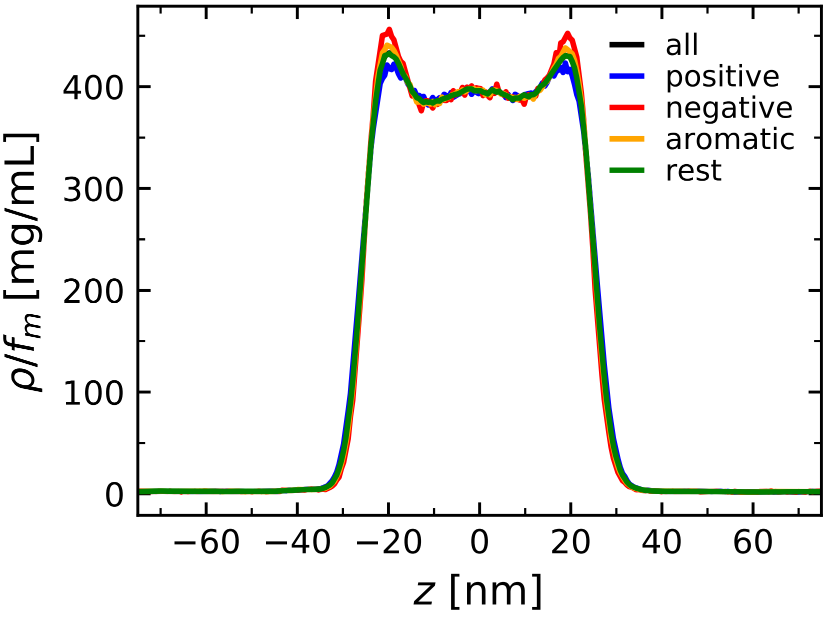


Figure S3. Density profile of A1-LCD (WT) simulated at $T = 280\,\mathrm{K}$, separated by residue type: Aromatic (F and Y, orange), positively charged (R and K, blue) and negatively charged (D, red). Each curve has been divided by the mass fraction of corresponding residues in the sequence, $f_m$.

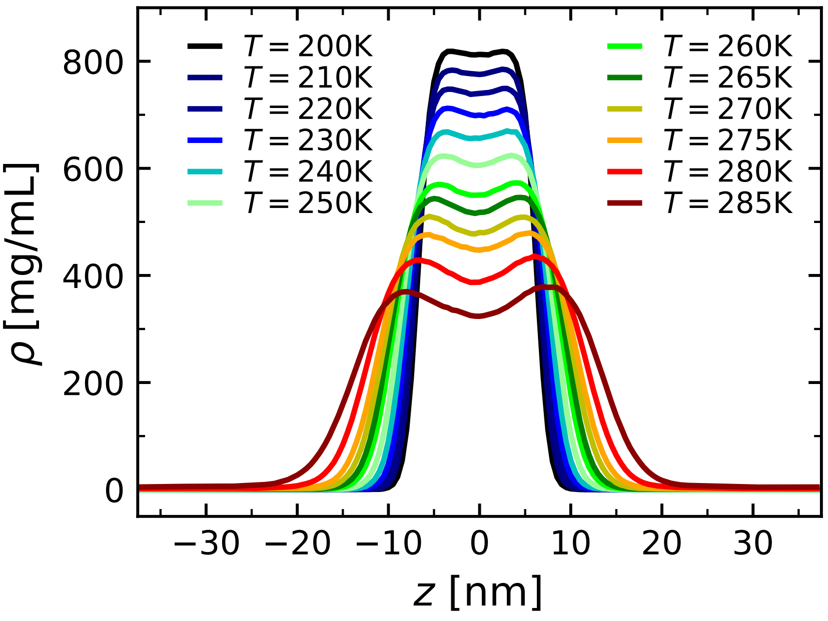


Figure S4. Density profiles obtained by simulations of 110 chains of A1-LCD (WT).

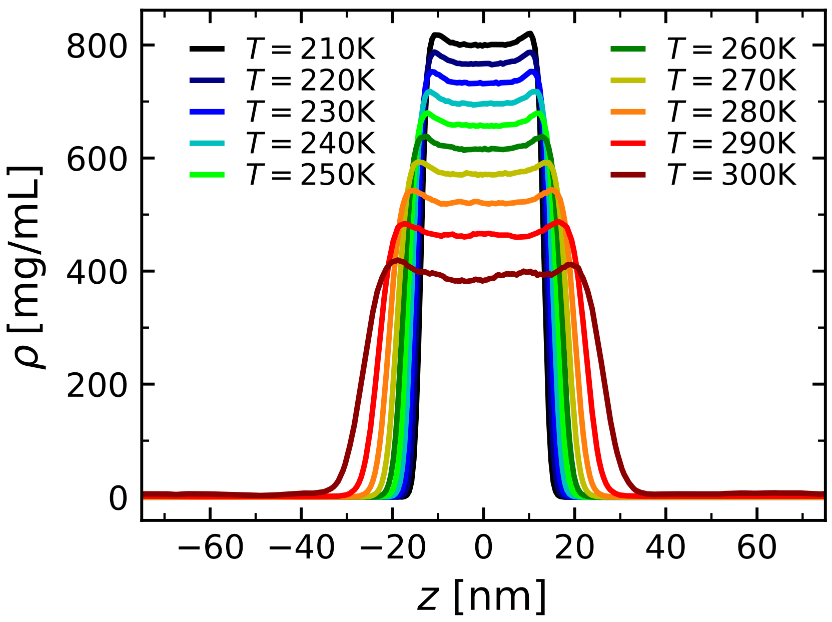


Figure S5. Density profiles obtained by simulations of 220 chains of A1-LCD ($Aro^+$).

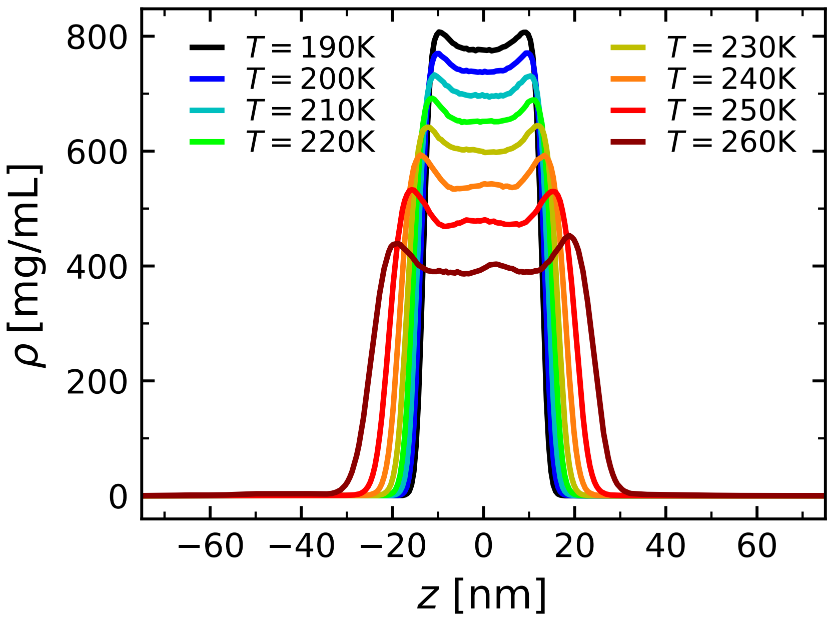


Figure S6. Density profiles obtained by simulations of 220 chains of A1-LCD ($Aro^-$).

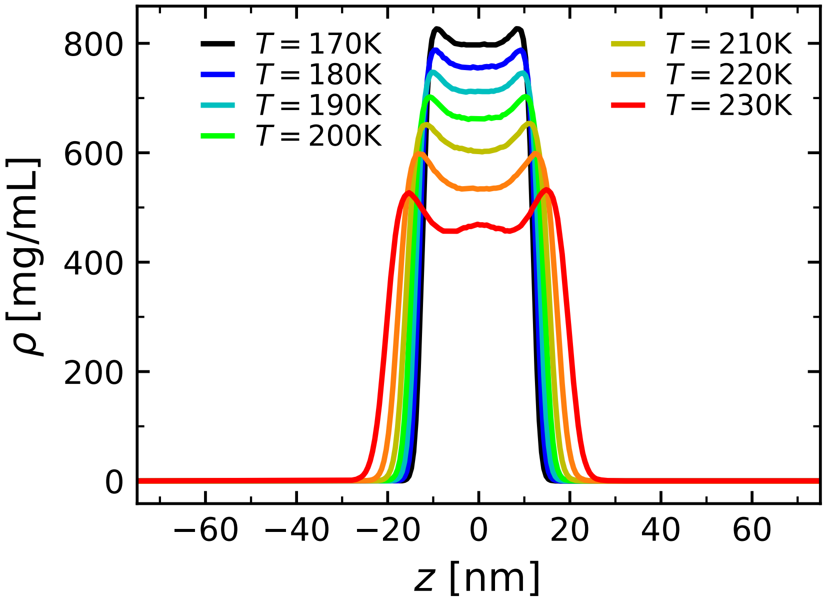


Figure S7. Density profiles obtained by simulations of 220 chains of A1-LCD (Aro$^{--}$).

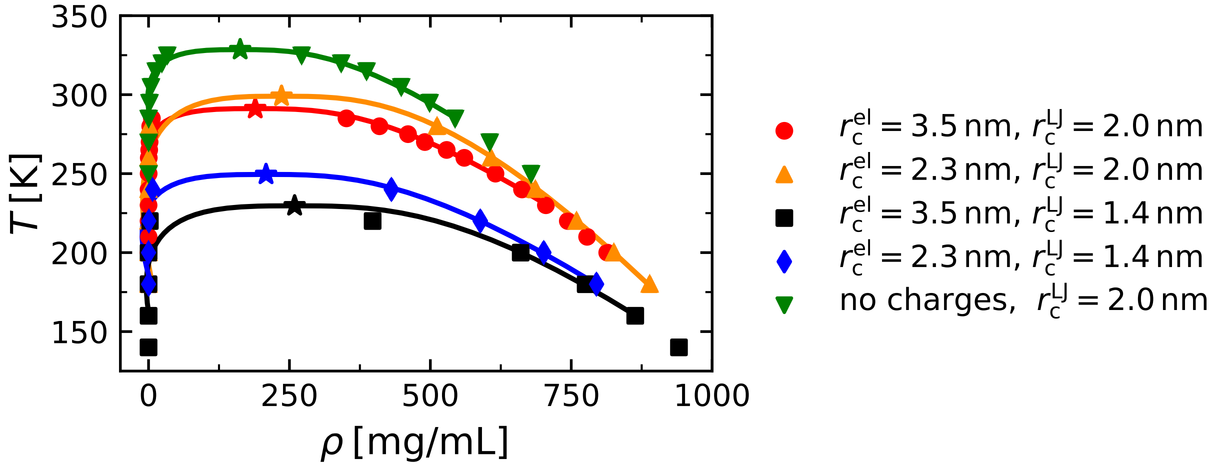


Figure S8. Phase diagrams of A1-LCD (WT) from simulations of 110 chains with different choices of cutoff radii, as indicated. Solid lines represent fits according to Eqs. (8) and (9) in the main text. Critical points are indicated by stars.

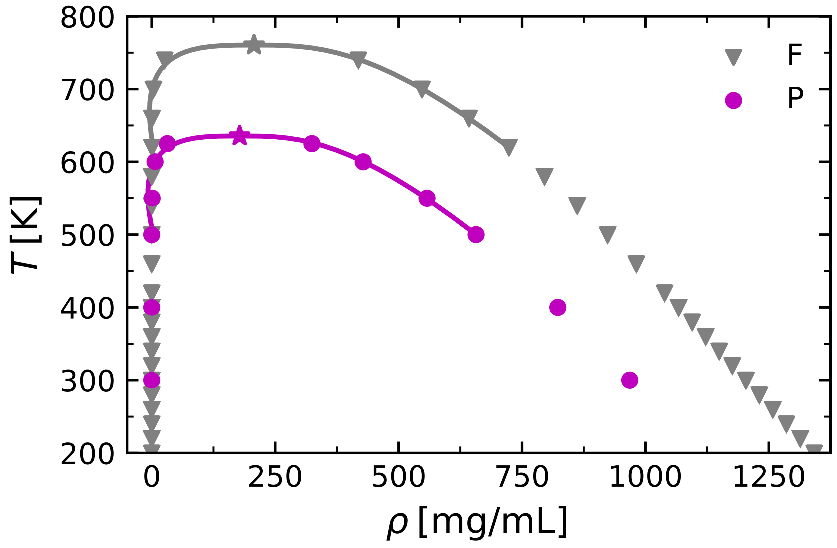


Figure S9. Phase diagrams of Phenylalanine (F) and Proline (P) homopolymers.

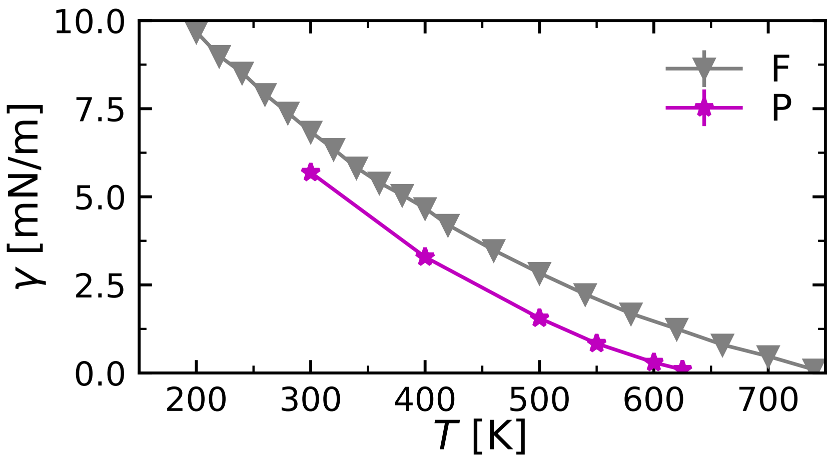


Figure S10. Surface tension $\gamma$ of Phenylalanine (F) and Proline (P) homopolymers as a function of $T$.
